## Supplementary material for "*SCAR-6* elncRNA locus epigenetically regulates *PROZ* and modulates coagulation and vascular function": Supply

—--------x—-----------------------x------------------------x—----------------------------x—---

### Supplementary Data

#### CRISPR-CAS9 mediated *scar-6* mutant generation

We designed the sgRNA targeting the 1st and 2nd exon of scar-6 lncRNA and checked the off targets of these sgRNA with 4-bp mismatch using Cas-OFFinder web tool (http://www.rgenome.net/cas-offinder/). sgRNAs showing no off- target with 4-bp mismatch and was selected. sgRNA IVT was generated as previously described in Varshney et al. 2015. The sgRNA an Cas9 RNP complex was injected into single cell zebrafish. The injected $F_0$ animals were then screened for any cardiovascular phenotype such as heart defect, vascular abnormality or hemmorage. The sgRNA targeting the 5' Region of the scar-6 did not show any cardiovascular related phenotype whereas the sgRNA targeting 3' region showed ~30% animals with hemmorage phenotype. We next followed these $F_0$ animals and they were grown to adulthood and then mutation screeneing for hetrozygous using heteroduplex mobility shift assay (HMA) on fin-clipped DNA was performed. In this assay, two PCR strands with different sequences (WT and mutant) are allowed to anneal, forming heteroduplexes. The mobility of these heteroduplex structures through the gel is influenced by their size, shape, and charge. By assessing the mobility shift of the heteroduplexes compared to the wildtype PCR template strands we can screen for the hetrozygous animals. We observed multiple bands in the PAGE gel compared to wild type, conforming mutation in the zebrafish. We observed multiple indels at the target loci in the $F_{0.}$ The $F_0$ zebrafish, positive for hetrozygous mutation was outcrossed with wild-type *gib004Tg*(*fli1: EGFP; gata1a: dsRed*) animals, and similarly, $F_1$ hetrozygous mutation positive animals were again outcrossed with wild-type *gib004Tg*(*fli1: EGFP; gata1a: dsRed*) animals to get a stable mutant line (Fig EV4F). $F_2$ mutant were genotyped and heterozygous mutant zebrafish animals were identified. We generated a stable mutant line with 12 bp deletion at the target loci of *scar-6* which were named *scar-6*$^{gib007\Delta12.}$ (Fig 3B)

**Table S1: Publically available datasets used in this study after reanalysis.**

| Sno. | Public data repository | Comment |
|---|---|---|
| 1 | GSE32900 | Development stages of zebrafish |
| 2 | GSE134055 | Tissues of zebrafish |
| 3 | PRJNA504385 | Zebrafish Endothelial cell |
| 4 | GSE133437 | CTCF ChIP-seq |

**Table S2 Details of prioritized 10 *scar* lncRNA genes**

| ***Name*** | **ZFLNC ID** | **Neighboring protein coding gene** |
|---|---|---|
| *scar-1* | ZFLNCG01330 | *ghrhra; adcyap1r1a; Scn12aa* |
| *scar-2* | ZFLNCG05739 | *prlh; rpe; ackr3a; tyw5; maip1* |
| *scar-3* | ZFLNCG09043 | *Apoa.4; Apoeb.2; Apoc* |
| *scar-4* | ZFLNCG09044 | *Apoa.4; Apoeb.2; Apoc* |
| *scar-5* | ZFLNCG11113 | *selp; sele* |
| *scar-6* | ZFLNCG00003 | *f7i; f10; prozb; pcid2; cul4a* |
| *scar-7* | ZFLNCG00988 | *myh7l; myh7* |
| *scar-8* | ZFLNCG00989 | *myh7l; myh7* |
| *scar-9* | ZFLNCG13000 | *sulf1; csrnp1b* |

| | | |
|---|---|---|
| *scar-10* | ZFLNCG07667 | *pax2a; cuedc2; hif1an; wnt8b; scdp; dnajb12a; trmt2b* |

**Table S3 : Genotypic percentage of animals showing hemmorage phenotype in incrossed scar-6 mutants.**

| | No-phenotype | | Phenotype | |
|---|---|---|---|---|
| | N | % | N | % |
| **Total** | 285 | 100.0 | | |
| **WT** | 69 | 24.2 | 0 | 0 |
| **Heterozygous** | 132 | 46.3 | 0 | 0 |
| **Homozygous** | 14 | 4.9 | 70 | 24.6 |

**Table S4 Primer details.**

| S. No | Name | Sequence (5'-3') |
|---|---|---|
| 1 | 3'RACE scar-6 F | GCCAAAAGCCTGTAGCCATT |
| 2 | 3'RACE nested scar-6 F | AGTCCTCGAGACACCACTGACCTCCATAGT |
| 3 | T7+scar-6 full length R | TAATACGACTCACTATAGGCGCAAGGGGAAATACGGCG |
| 4 | scar-6 full length F | CGAAGCATTGATGAACCATTC |
| 5 | T7+scar-6 full length F | TAATACGACTCACTATAGGGCGAAGCATTGATGAACCATTC |
| 6 | scar-6 full length R | CGCAAGGGGAAATACGGCG |
| 7 | RT_zff10F | AGAAGAATGTGGTCTGCTCG |

| | | |
|---|---|---|
| 8 | RT_zff10R | ATGCTGTCGGTCTGGTTGTT |
| 9 | RT_zfprozbF | GGAGACCAGTGCAGATCTAA |
| 10 | RT_zfprozbR | CGTCGCTGTTGAGTGTGTAT |
| 11 | zf actinb F | AA TTGCTTCCGAGGCGcgctggagctaatcgatga |
| 12 | zf actinb R | GGTGGCTCCAACTCGgtgaccatgtggagtcagctt |
| 13 | human_scar-6-F | CTGCCCTCCGCGCAGCATGGA |
| 14 | human_scar-6-R | GGATCGACAGGTCCATGAAAAC |
| 15 | scar-6 exon2 F | TTATTGGTGTCTTGTGCCGC |
| 16 | scar-6 exon2 R | AAGGGGAAATACGGCGTCTA |
| 17 | PAR1(f2r) F | CCC CCG GCT AAA AAG ACT TA |
| 18 | PAR1(f2r) R | GGCTCCGTATATCCAGTTGT |
| 19 | PAR2a F (f2rl1.1) | ATTGAACAGCAAGCTCACGC |
| 20 | PAR2a R | CAGTACCTCTGGACGCTAAT |
| 21 | PAR2b F (f2rl1.2) | CATCTGGACGCCTCTAAAGA |
| 22 | PAR2b R | TGATGATGTTGACGTCGTGG |
| 23 | PAR3 F (f2rl2) | TCATCAATCACACCGCTGGA |
| 24 | PAR3 R | AGCTTGCAAGCTAGTTCACC |
| 25 | T7+f10 F | TAATACGACTCACTATAGGGTCC TGA ACT CTG CGA GAA TG |

| | | |
|---|---|---|
| 26 | T3+f10 R | GCAATTAACCCTCACTAAAGGGAAATTGCTGGACTCGATGC |
| 27 | T7+prozb F | TAATACGACTCACTATAGGGAAA TCC CAG TGT CCA TCT GC |
| 28 | T3+prozb R | GCAATTAACCCTCACTAAAGGGCCTGTCAGAAAGGCTGTTT |
| 29 | f10 F | TCC TGA ACT CTG CGA GAA TG |
| 30 | f10 R | GAAATTGCTGGACTCGATGC |
| 31 | prozb F | AAATCCCAGTGTCCATCTGC |
| 32 | prozb R | GCCTGTCAGAAAGGCTGTTT |
| 33 | CTCF_RT_six6a F | AATGACCGCTGACAATAGCG |
| 34 | CTCF_RT_six6a R | GAGCAGGGTTAAACACTGAC |
| 35 | CTCF_RT_scar-6 F | AATACATCATCCCCGCGT |
| 36 | CTCF_RT_scar-6 R | CGGTTAGTTTCTGCAGGATG |
| 37 | CTCF_RT_f7 F | CTACAGTGTGTATGCGGTGC |
| 38 | CTCF_RT_f7 R | GTATCCGGCGCAGAACAT |
| 39 | vcam1b F zf | GATGCTGGAACTTACCAGTG |
| 40 | vcam1b R zf | CTTGACTGTGGACTTGCTAC |

| | | |
|---|---|---|
| 41 | vcam1aF zf | CTGCTGATAATCTGGGCAAG |
| 42 | vcam1aR zf | TCACCAAGCTTTACTGAGGC |
| 43 | cdh5 zf F | GGA CAG AGA GCA AGA ATC CT |
| 44 | cdh5 zf R | CTCACAGACATACACGTAGC |
| 45 | claudin 11a zf F | ATCACTGCATCACACTCACG |
| 46 | claudin 11a zf R | GATCGCTTGTTCTTGGCACT |
| 47 | *claudin 5a zf F* | CCA GAT GCA ATG CAA AGT GC |
| 48 | *claudin 5a zf R* | AGTGGCACAAGCACGAAGAT |
| 49 | iNOS_zfF nos2a | CTGAACTGATCTTGGAGGTC |
| 50 | iNOS_zfRa | ACCACCAACTTCCATGAGCA |
| 51 | iNOS_zfF nos2b | GGAACTTCTGTGATACCCAG |
| 52 | iNOS_zfRb | TGTGCTGCATGAAGGACTCT |
| 53 | *Claudin-2F | CATGAAGGGTCTTTGGATGG |
| 54 | *Claudin-2R | ACTGAAGATCCGTCCATGCA |
| 55 | E-selectinF | GCAGCTTCAGTTGTGCAGAA |
| 56 | E-selectinR | CATTACTGTCAAGCTGGTGG |
| 57 | ICAM-1 F | GAGACGAAGGACCTCAACAT |
| 58 | ICAM-1 R | GGTTTGCCTTCACTGTACAG |
| 59 | T7+sgRNA scar-6 | TAATACGACTCACTATAGAAATACGGCGTCTACACACGTTT |

| | | |
|---|---|---|
| | | TAGAGCTAGAAATAGC |
| 60 | scar-6 CR F | CCAGGAGGAGAAAGATGCAT |
| 61 | scar-6 CR R | TTATTGGTGTCTTGTGCCGC |
| 62 | BS-scar-6 F | TGGTGATATATTAAATATATTTTTTATGTT |
| 63 | BS-scar-6 R | CTTTCCTCAAAAAAATCAAACTCTT |
| 64 | prdm14 chip F_scar-6 | AGGTGTGGTGAGCTGGGG |
| 65 | prdm14 chip R_scar-6 | GTCTCTGGCATGACTTTCGT |
| 66 | prozb full length R | TCAGGGTTGCTCCCTCTCGG |
| 67 | prozb full length F +t7 | TAATACGACTCACTATAGGATGGAGTCGCTTGTATATCG |
| 68 | scar-6 morpholino | GCATTTAATTACTCAGTCCACTAACAGC |

**Supplymentary Figures and Legends**

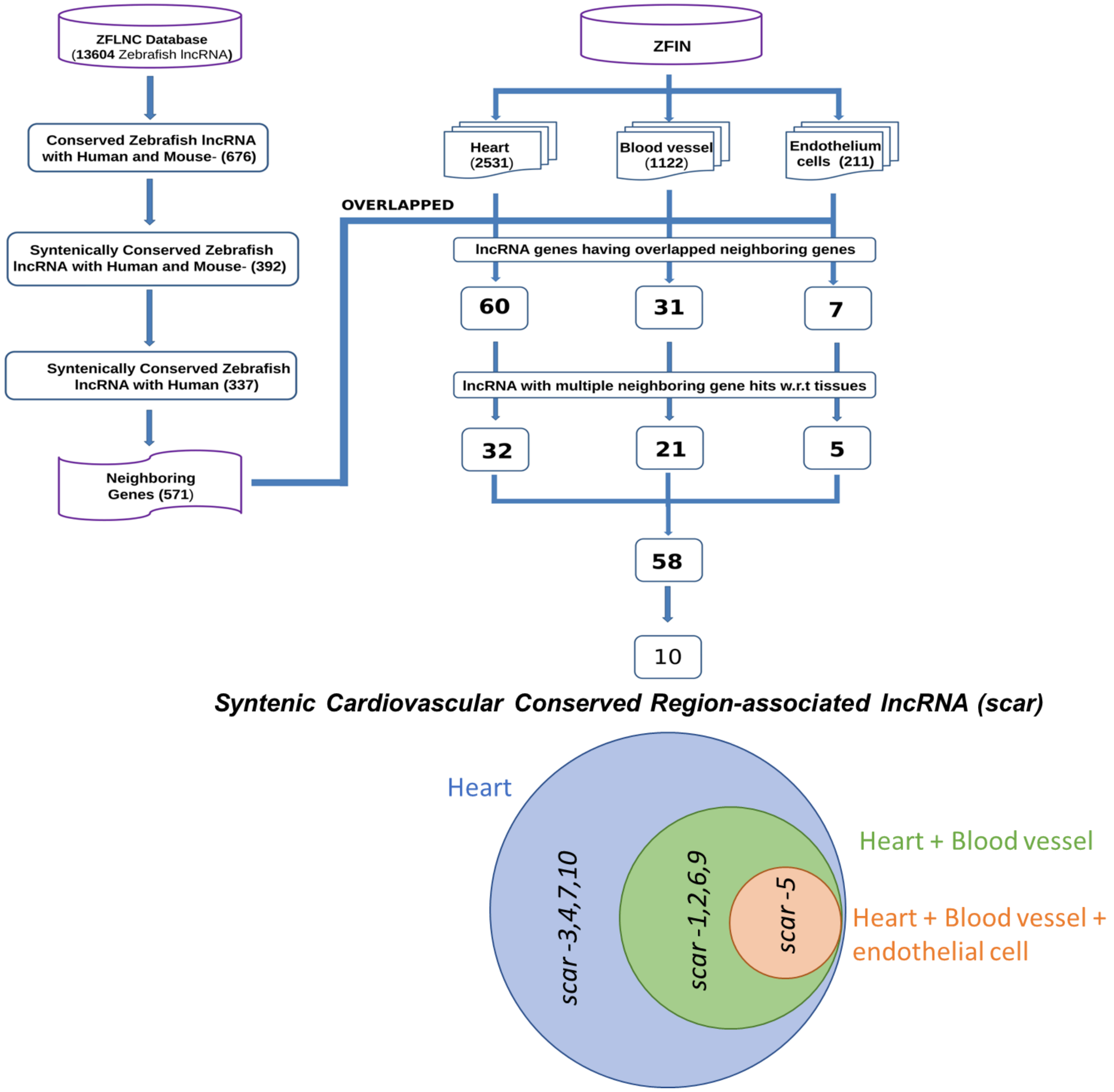


**Supplementary Figure 1: -** Schematic of pipeline used for subset selection of syntenic lncRNAs with its neighbouring protein-coding genes associated in the cardiovascular system.

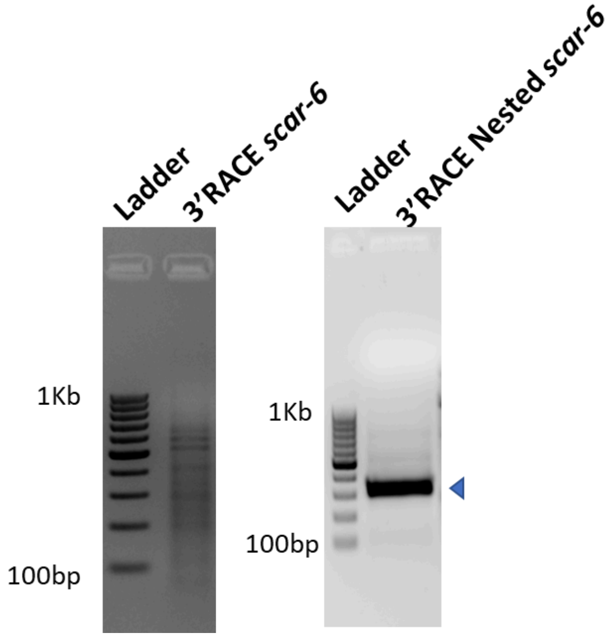


**Supplementary Figure 2: -** Agarose gel image for 3'-RACE of *scar-6* lncRNA followed by nested PCR resulted in expected size of the transcript.

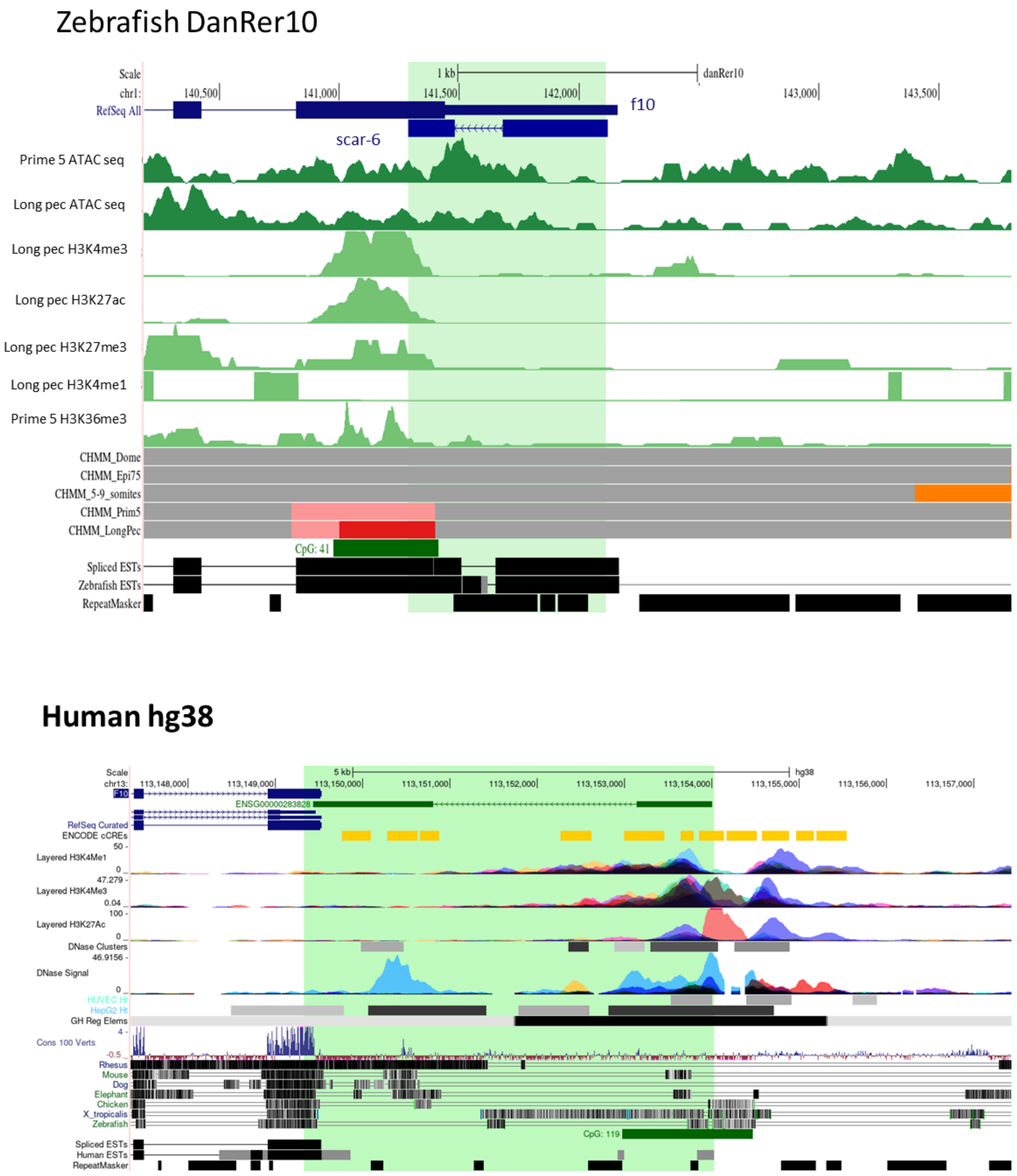


**Supplementary Figure 3: -** Genomic context of *scar-6* locus in zerbrafish and Human with H3K4me1, H3K4me3, H3K27ac, H3K36me3, H3K27me3, chromatin state marks (ENCODE Project Consortium *et al*, 2020; Baranasic *et al*, 2022)

```
ENST00000639766.1      TTCTGGGAGGTCGAACACCTGTCCACTTTGATTTTCCCTGACTTCGCAACATCTGTTTAC
ZFLNCT00004            --------------------------------------------CGAAGC----ATTGAT
                                                                   ** * *     ** *

ENST00000639766.1      GAACGGTTTTGCCCAGCTCATGACTTTTTTGGGGAGTTTTTGTACTAGAACTGTT--GAA
ZFLNCT00004            GAACCATTCTACCTCCATAATGT-------------GATTTTACTAGAAATACCATGAT
                       ****  ** * **    * ***                ** ******** *     **

ENST00000639766.1      CTT--TGCTTAGGGACATTACACAAGACGCAAACTGCAGGA------------------
ZFLNCT00004            ATTAATACACATGAGGATTACACTAAAGCCACAGTTCACCCAAAAATCTACATTTACCCA
                        **  * *  * *   ******* * *  ** * * **

ENST00000639766.1      -----------------------------------------------------GACGTTT
ZFLNCT00004            CTATTTACTGCCAAACATTTAAGAGTTTCTTTCTTCTTTTAAACACAAAAGAAAATATTC
                                                                          *  **

ENST00000639766.1      GGGA----G---ACACACGGCAGGCCTTGTCCGGAATCTTTTCTGTTTA-----------
ZFLNCT00004            TAAAGAAAGCCAAAAGCCTGTAGCCATTGTAATGAAA----TCCGATTAAAATTAAAATA
                          *    *   * *  * * ** * ****   ***     ** * ***

ENST00000639766.1      ------CCGAAAGTTCATTTGATTTTCATGGCCAGTTAAC-TTGAGGCC---TAGGGACA
ZFLNCT00004            TTCATAATTAAATTGCAACTTGTTATCATTAATTATTAATATTCATACTTTTAAGTTACA
                               *** * **  *  ** ****      ****  ** *  *      **  ***

ENST00000639766.1      GTGCCGGATTGCAGCAGGGCAAGGACTAAACATCTGGACCCTAGGAGAGGAGGAGCCCCC
ZFLNCT00004            CCACTGACCT---CCATAGTAAG-AATAAAAAC---------------------------
                          * *   *    **  * *** * **** *

ENST00000639766.1      CGAAGCCCCGCCTGCCACGCCACCACCACTCAGCACCTG-AGGAGCTCCAGAAGCTTCCA
ZFLNCT00004            ------------------------------CAATAGATAAAGGTGTTC-AACATTCTTCA
                                                     **  *  *  *** * ** *  *   * **

ENST00000639766.1      GAAGCTCCTGGCCAGCTTCACTCCAGAGAC--------TGATTTCAGTGGCCAG--GCTG
ZFLNCT00004            GAACTTCTTCTTTTGTTTTATAACAGAAAACTAGAAGCTGGTAA--AGGACGAGTAACTT
                       ***  ** *     * ** *   **** *          ** *      * * **   **

ENST00000639766.1      TGGCCTGGATGCAGGCATGC-TAAAGCCCACAGGGGGTTCTAAGGGTGGCCACGGTGGGG
ZFLNCT00004            ----TTGGGTGAACTCTCCCTTAAAG--CACAGGTAATTAAATGCATATGAA--------
                            *** ** *  *    * *****  ******   **  * *  *    *

ENST00000639766.1      GCCAACTGTGCCAGGTCAGGTCGCTGCCTGGTGAGCAGGTCACCTGCCCGGGGGATGCCT
ZFLNCT00004            ------------------------------------------------------------

ENST00000639766.1      GCCAACTGTGCCAGGTCAGGTCGCTGCCTGGTGAGCAGGTCACCTGCCCGGGGGATGCCT
ZFLNCT00004            ------------------------------------------------------------

ENST00000639766.1      GTGGCCCCCACCACACAAGCCTGAGCTGGGGTGGGGCCTGGCACCGTGCAGTTGAAGAGC
ZFLNCT00004            ------------------------------------------------------------

ENST00000639766.1      ACCTGGTTGTTTAAGTTAATGTAAAATGTCCGCCCCAATGAGAAGCTTTCTGGACGTTTC
ZFLNCT00004            ------------------------------------------------------------

ENST00000639766.1      CCTCCCAGGGAGTCTGCGGGTCATAGGCAGTCACCGGGGAGGTGTCCAATTCCCAAGAGG
ZFLNCT00004            --------------------------------------AGATGAGGCTTTCCTCAGAGG
                                                             ** **      ****  *****

ENST00000639766.1      AACAAGAGAGACCAGGCCAAGTGGGGGGTCTTGGG--GAGGCTGCGCCAACGTGTGGGCC
ZFLNCT00004            AGTCAG----ACTCTTCTTATTGG--TGTCTTGTGCCGCAGCTCTCGCTTTGCTTTGGGT
                       *   **    **    *  * ***   ****** *  *  ***    *    *  * **

ENST00000639766.1      CCTCCCGACGCCACGGGGCCTGCACAGGCTCCCATCTGCCTCTGGGC----GTGACCCAG
ZFLNCT00004            TTTAAAGACGCTCCGGTCTCTGGCATGACTTTC------GTCATTGCGTTATTGATCCAC
                         *   *****  ***   ***    * **  *       **   **      *** ***

ENST00000639766.1      ATGCAGGAAGGTATCTGGGGAAAGGAATGCCCATGGCAGTCATGGCACAGTATCATCCCT
ZFLNCT00004            ATG-----ATGTATTTGGAGACCTGTGTGT------------AGACGCCGTATTTCCCCT
                       ***     * **** *** **   *  **               * * * ****   ****

ENST00000639766.1      GGAGCCTCCATCCTGCCCCCACAGACTAGAATATTCCCTGGAGGAATCCCCAGCCTAGCT
ZFLNCT00004            TGCG--------------------------------------------------------
                        * *

ENST00000639766.1      GGGAAGGGCGGAAGCTGGTGTTCCAGACAGACACTTCACCTTCCCACCTCAACAGGGCGG
ZFLNCT00004            ------------------------------------------------------------

ENST00000639766.1      GAGAAGCCAGGGCAAGGCGGCAATCGAGAGACAAACCAGGCCTTGAGTGGGATCTCACTT
ZFLNCT00004            ------------------------------------------------------------
```

**Supplementary Figure 4: -** ClusterW based sequence alignment of zebrafish *scar-6 (ZFLNCT00004)* and human *SCAR-6 (ENST00000639766.1)* lncRNA transcript.

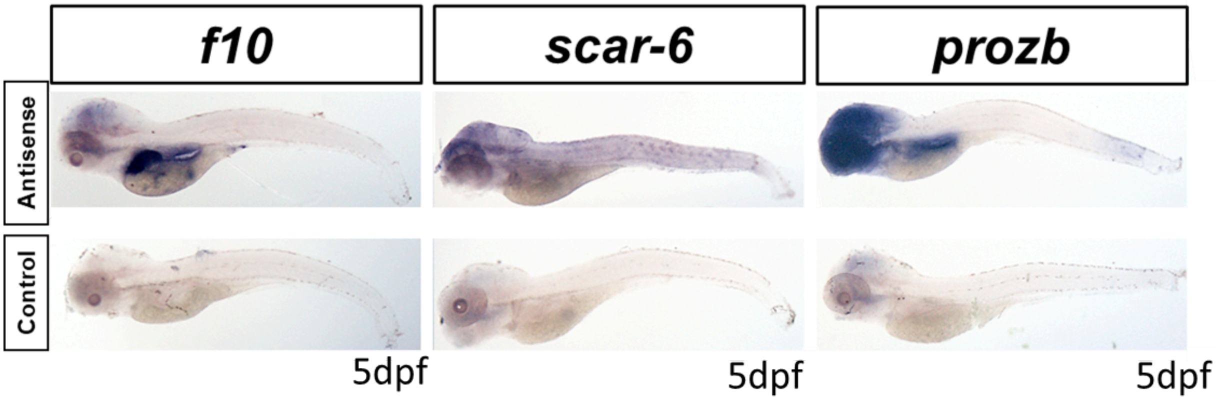


**Supplementary Figures 5: -**Whole mount in-situ hybridization expression analysis of *scar-6, f10,* and *prozb* transcripts of zebrafish in 5dpf embryo. Sense probes were used as controls for *f10* and *prozb*. No probe control was used for *scar-6.*

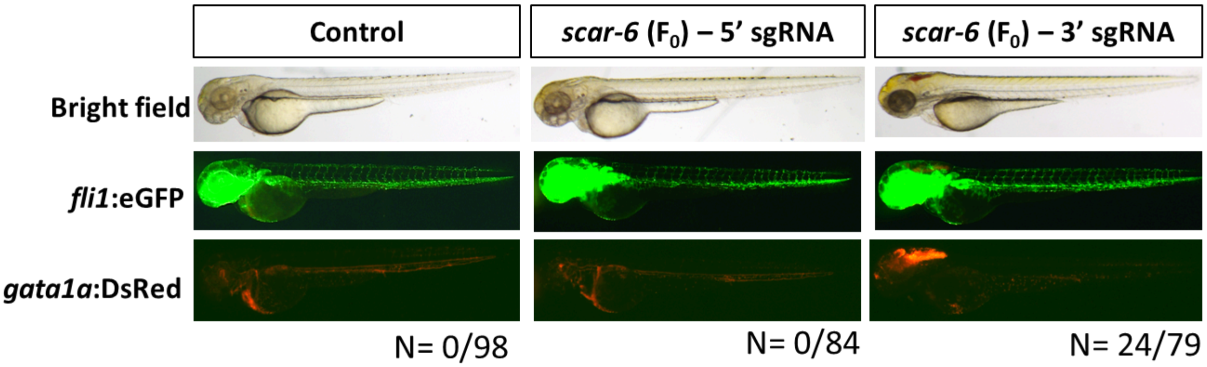


**Supplementary Figures 6:** -Representative image showing transgenic gib004Tg(*fli1a:*EGFP;*gata1a*:DsRed) 3dpf zebrafish injected with RNP complex of CRISPR-Cas9 targeting 5' and 3' region of *scar-6* gene (5x resolution)

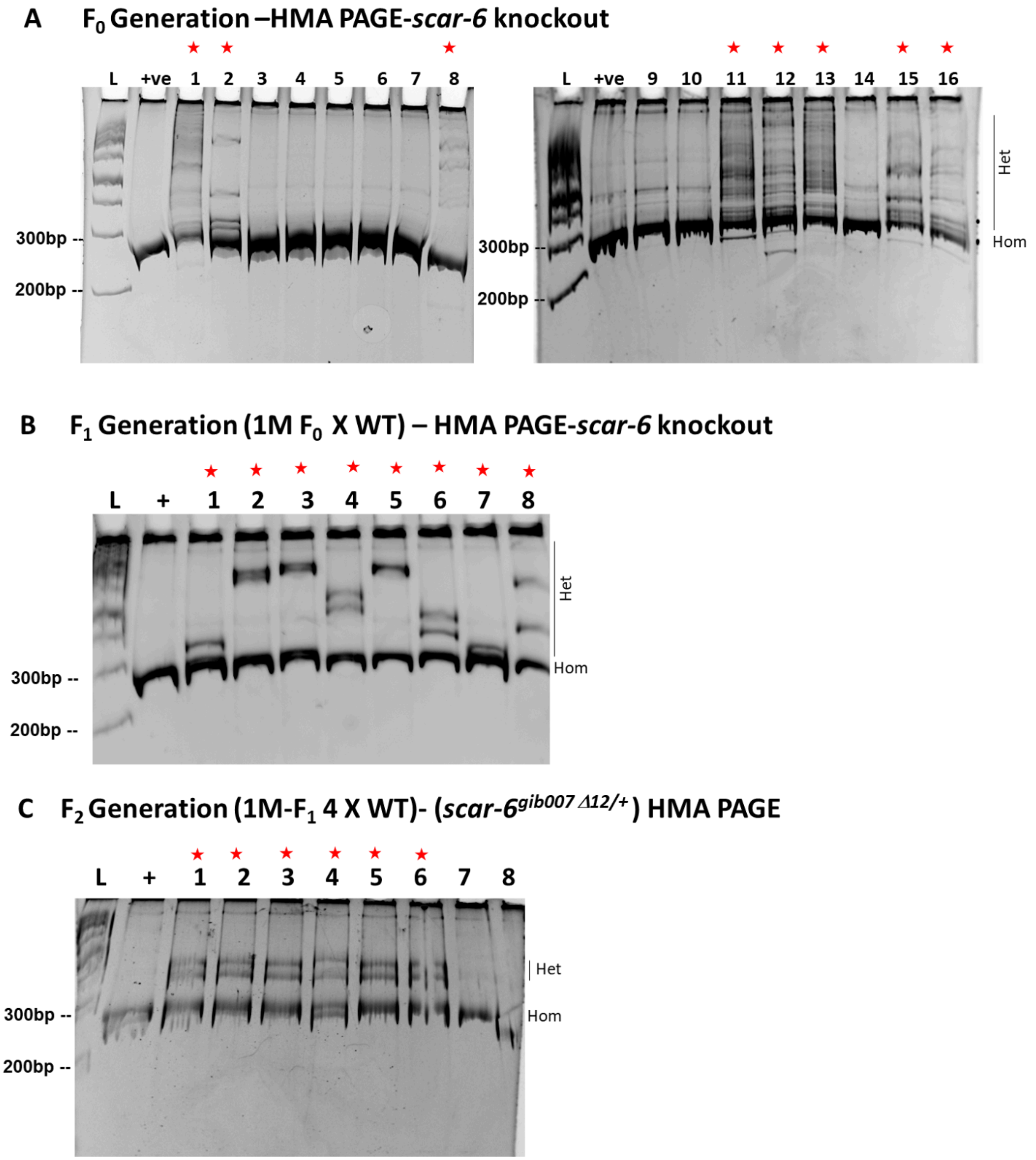


**Supplementary Figures 7: - CRISPR-Cas9 mediated mutant of *scar-6* lncRNA**

[A] Heteroduplex mobility assay (HMA) PAGE gel for *scar-6* target region in $F_0$ zebrafish.

[B] HMA-PAGE gel for *scar-6* target region in $F_1$ zebrafish.(star represent positive for mutation)

[D] HMA-PAGE gel for *scar-6* target region in $F_2$ zebrafish.(star represent positive for mutation).; Het- Heteroduplex, Hom- Homoduplex.

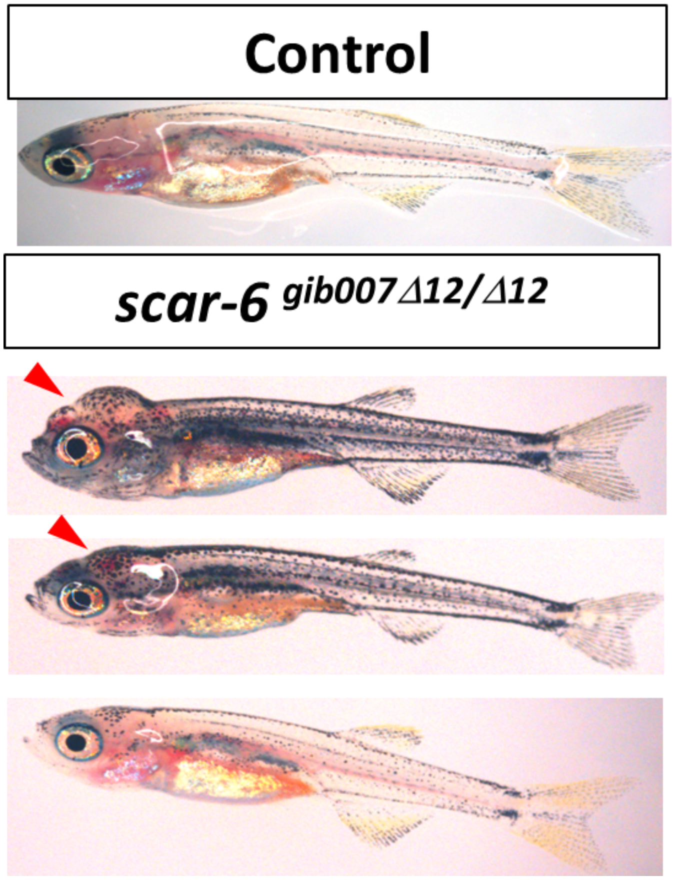


**Supplementary Figures 8: -** Raw image of 30 dpf zebrafish from wild type and *scar-6*$^{gib007\Delta12/\Delta12}$ animals which were closely monitored for their survival.

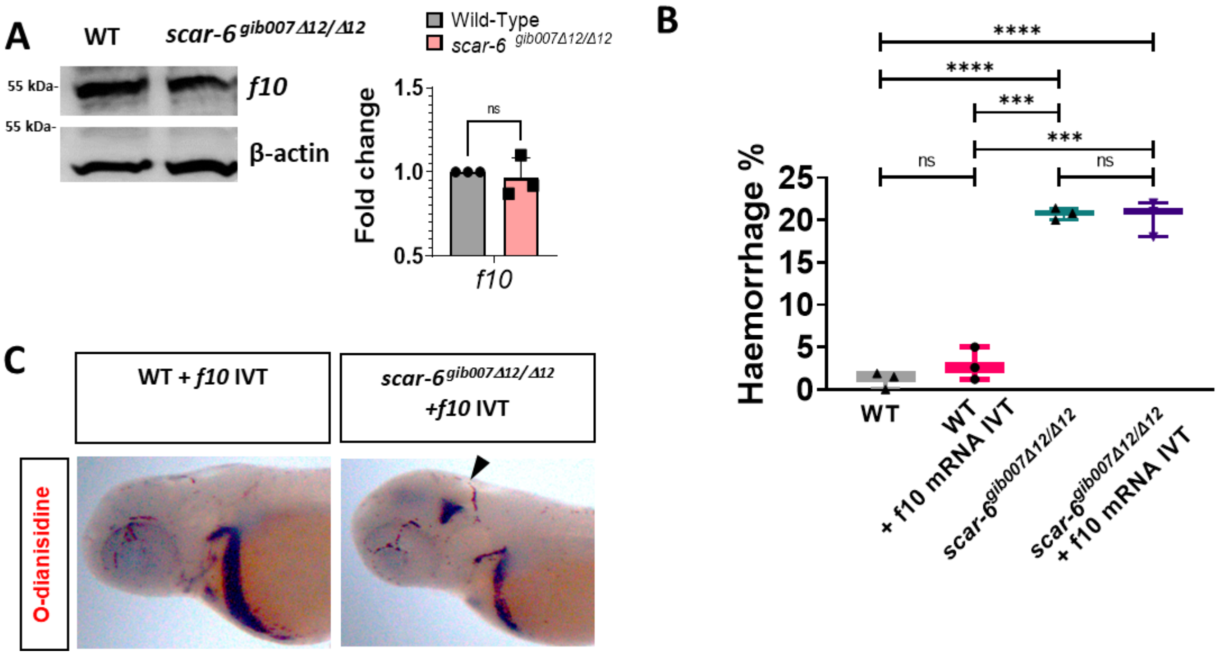


**Supplementary Figures 9: Rescue of *scar-6* mutant with *f10* mRNA**

**[A]**Western blot of *f10* in wild type and *scar-6*$^{gib007\Delta12/\Delta12}$ mutant zebrafish. The bar plot represents the quantification of the western blot from 3 independent biological replicates plotted as mean fold change ± standard deviation.ns- not significant (two-tailed unpaired t-test)

**[B]**Box plot representing the percentage of animals exhibiting haemorrhage phenotype in wild type(WT), WT injected with f10 IVT RNA (100 ng/uL), *scar-6*$^{gib007\Delta12/\Delta12}$ mutant and *scar-6*$^{gib007\Delta12/\Delta12}$ mutant zebrafish injected with f10 IVT RNA (100 ng/uL). Data from 3 independent biological replicates plotted as mean percentage ± standard deviation; *** $p<0.001$, **** $p<0.0001$ (two-tailed unpaired t-test).

**[C]**Representative image showing the cranial region of 3dpf zebrafish with o-dianisidine staining of RBC blood cells in wild type and *scar-6*$^{gib007\Delta12/\Delta12}$ mutant zebrafish injected with f10 IVT RNA (100 ng/uL). (4x magnification)

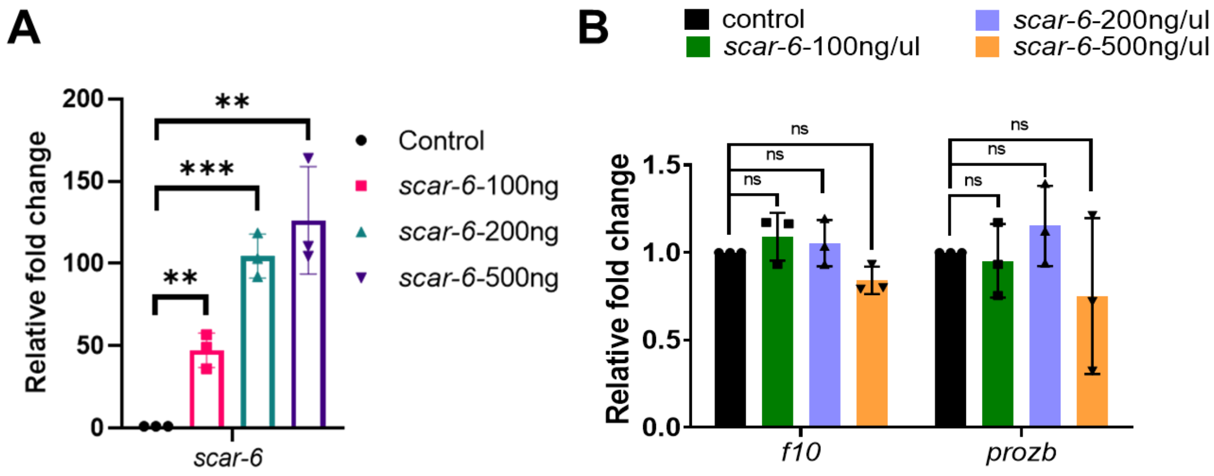


**Supplementary Figures 10: -Overexpression of scar-6 lncRNA in zebrafish**

**[A]** Relative fold change expression of *scar-6* at 3dpf upon *scar-6* IVT RNA overexpression in zebrafish at different concentrations (100, 200 500 ng/uL). Data from 3 independent biological replicates plotted as mean fold change ± standard deviation. ** P<0.01, *** p<0.001, **** p<0.0001 (two-tailed unpaired t-test).

**[B]** Relative fold change expression of *f10,* and *prozb* upon *scar-6* IVT RNA overexpression in zebrafish at different concentration (100, 200 500 ng/uL). Data from 3 independent biological replicates plotted as mean fold change ± standard deviation. ns - not significant (two-tailed unpaired t-test).

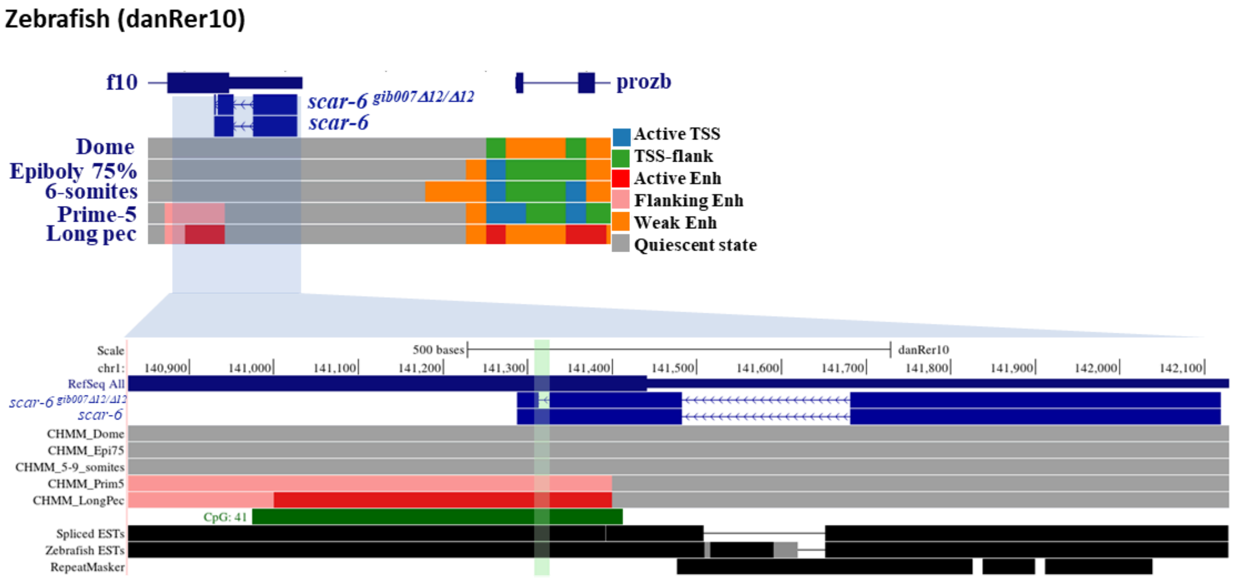


**Supplementary Figures 11:-** UCSC genome browser snapshot of zebrafish *scar-6* locus with chromatin annotation marks from danio-code data across 5 developmental stages.

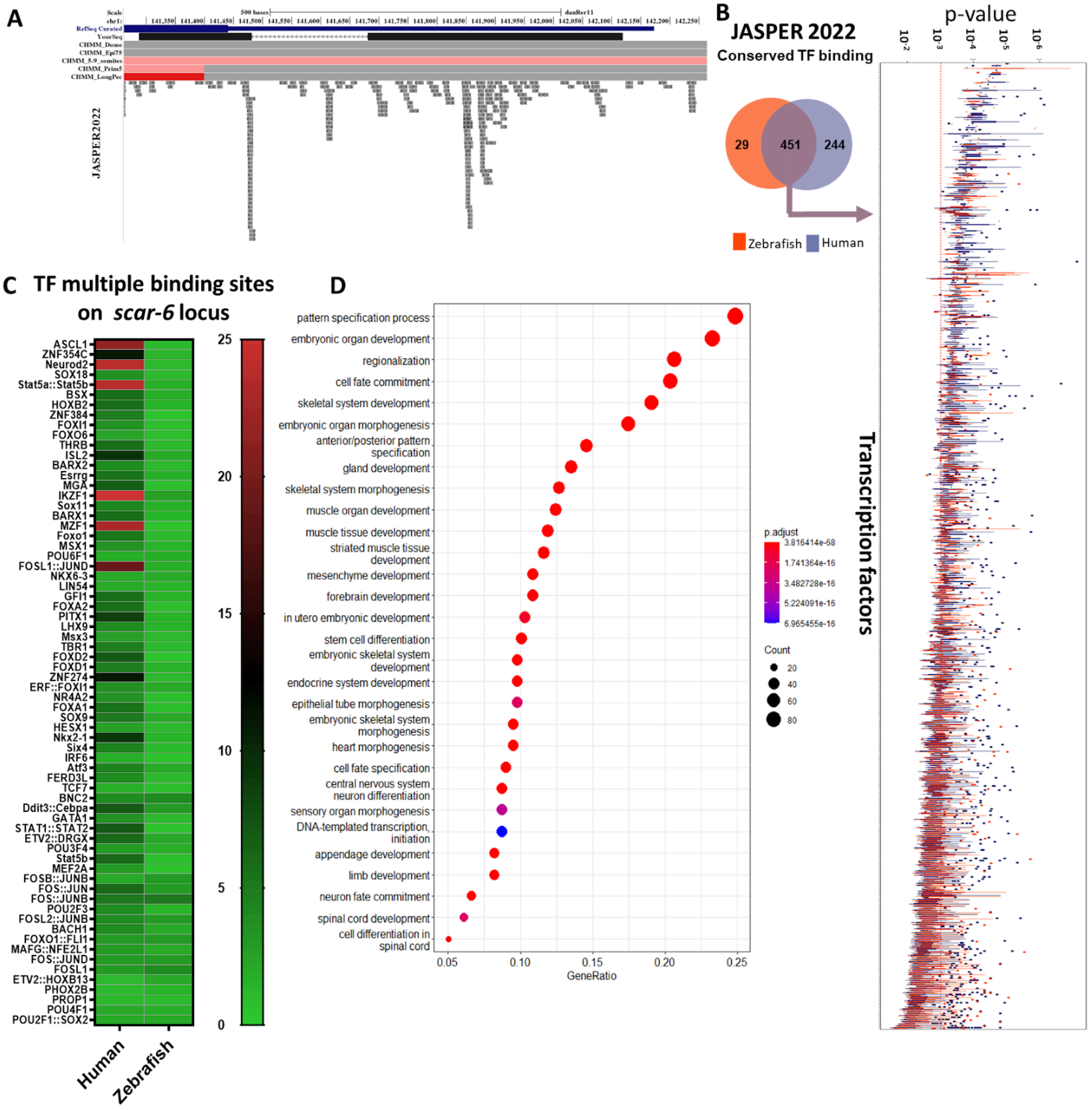


**Supplementary Figures 12: - Cis-Regulatory feature of *scar-6* locus**

[A]UCSC genome browser screenshot of *scar-6* locus and enrichment of transcription factors (TF) from JASPER 2022 data set with p value <0.001.

[B] Overlap of TF from *SCAR-6* locus of human and *scar-6* locus of zebrafish extracted from JASPER 2022 database. Box plot representing TF and their motif p-value. The red line indicate p value=0.001

[C]Heatmap representing the top 64 TF which showed multiple binding sites on *SCAR-6* locus of human and *scar-6* locus of zebrafish.

[D]Biological Gene ontology for common TF between human and zebrafish *scar-6* lncRNA locus.

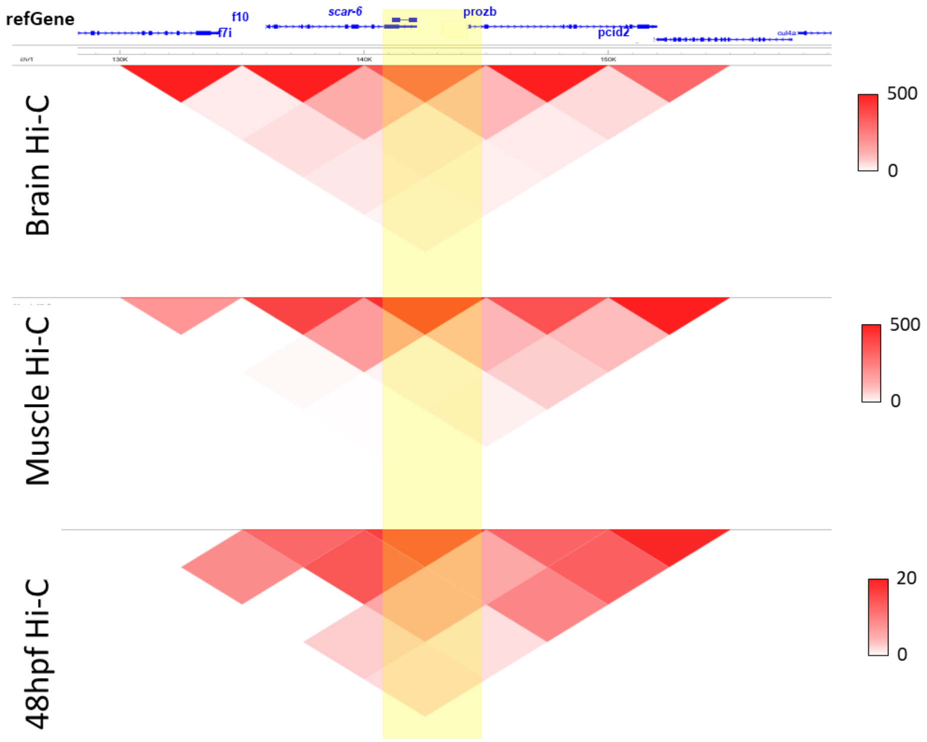


**Supplementary Figures 13:-** Hi-C heatmap representation of zebrafish *scar-6* locus in brain, muscle and 48 hpf zebrafish at 5kb resolution (Yang *et al,* 2020, Franke, M *et al*. 2020).

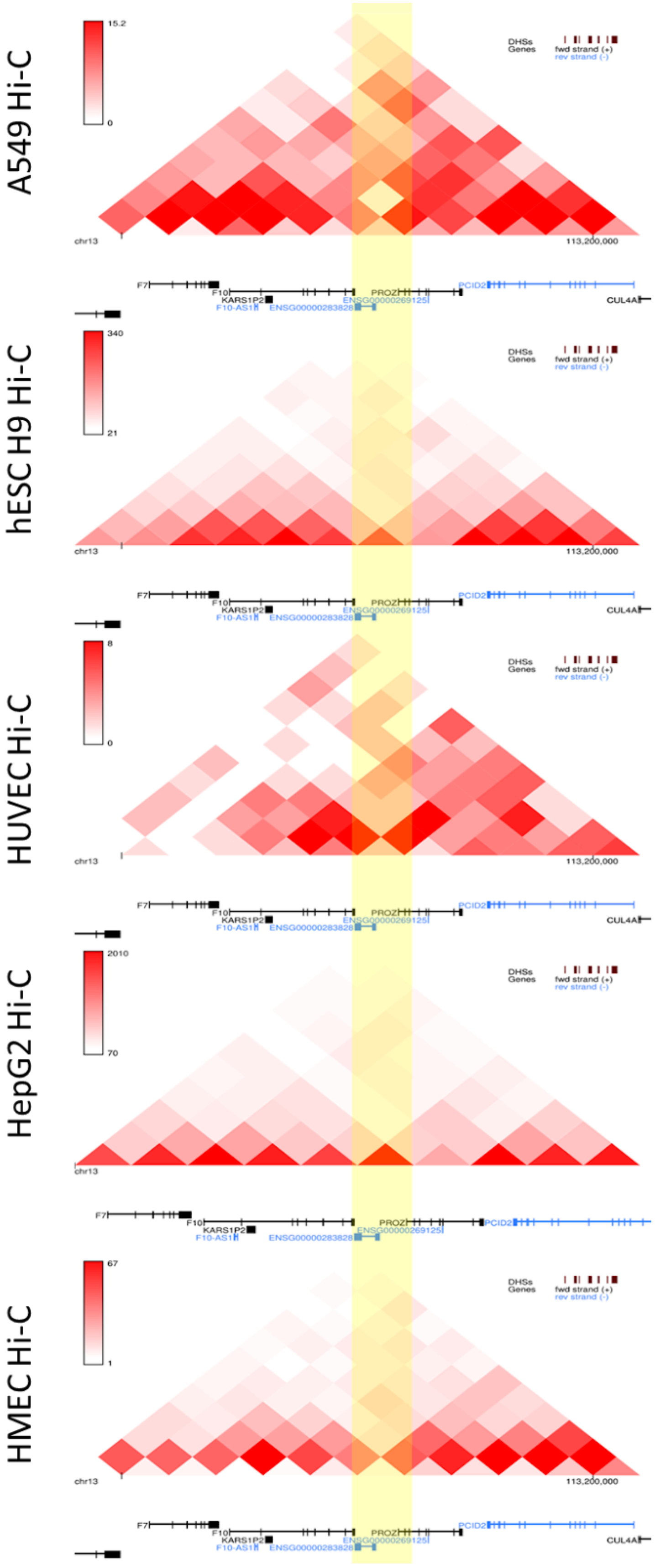
A549 Hi-C
15.2
0
hESC H9 Hi-C
340
21
HUVEC Hi-C
8
0
HepG2 Hi-C
2010
70
HMEC Hi-C
67
1
DHSs
Genes
fwd strand (+)
rev strand (-)
chr13
113,200,000
F7
F10
KARS1P2
F10-AS1
ENSG00000283828
ENSG00000269125
PROZ
PCID2
CUL4A

**Supplementary Figures 14: -** Hi-C heatmap representation of Human *SCAR-6* locus in A549, hESC-H9, HUVEC, HepG2 and HMEC at 10kb resolution from ENCODE database (ENCODE Project Consortium *et al*, 2020; Wang *et al*, 2018)

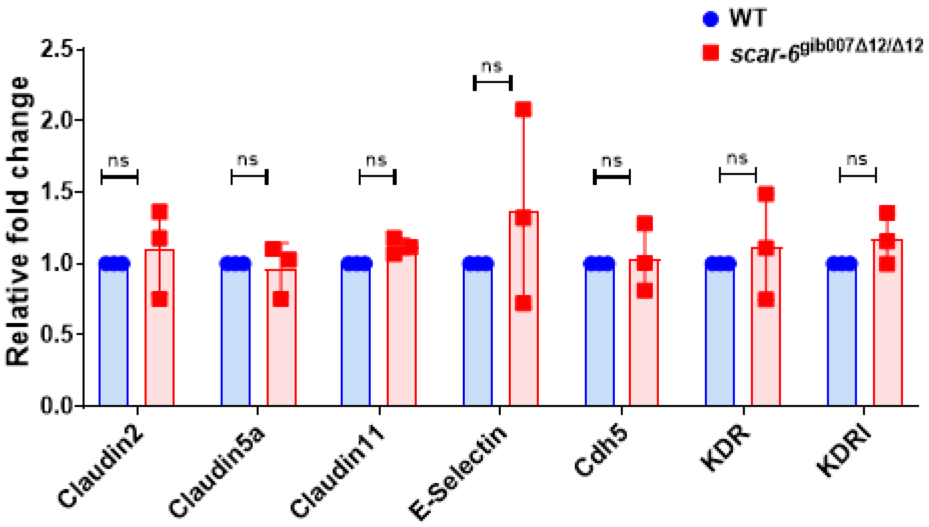


**Supplementary Figures 15: -** qRT-PCR of the endothelial associated gene downstream of Nf-kB. when compared between wild type and *scar-6*$^{gib007\Delta12/\Delta12}$ mutant zebrafish. Data from 3 independent biological replicates plotted as mean fold change ± standard deviation; ns- not significant (two-tailed unpaired t-test).

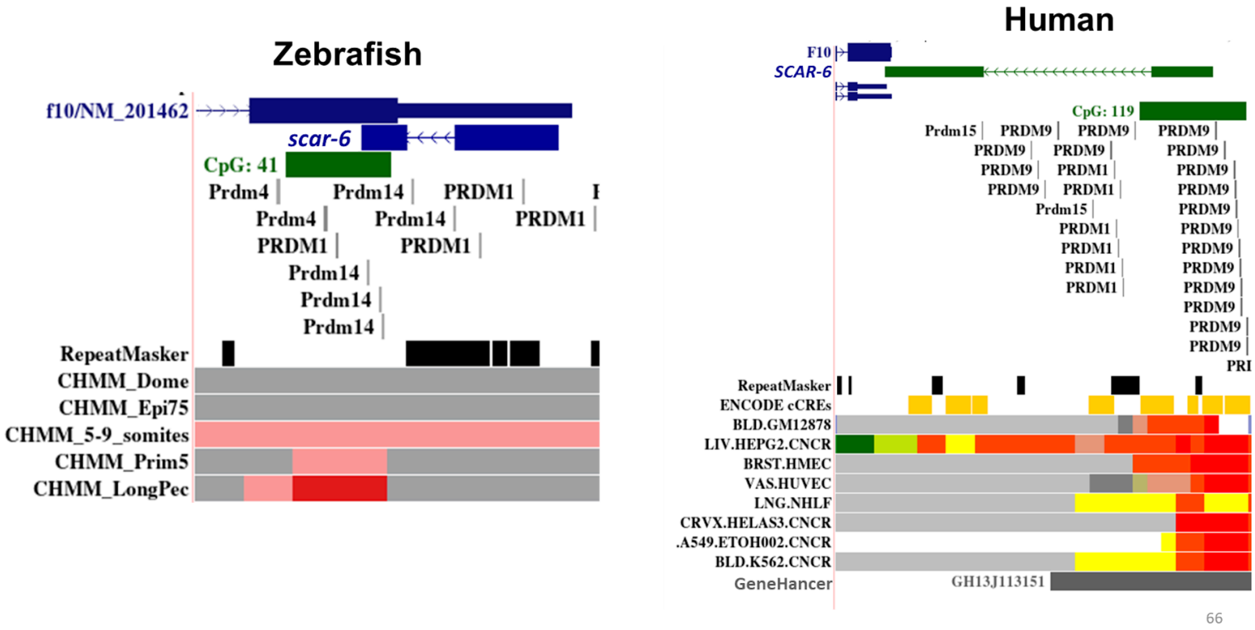


**Supplementary Figures 16 : -** UCSC genome browser screenshot of zebrafish *scar-6* and human *SCAR-6* locus depicting TF binding motifs of *PRDM* family proteins from JASPER 2022 database.

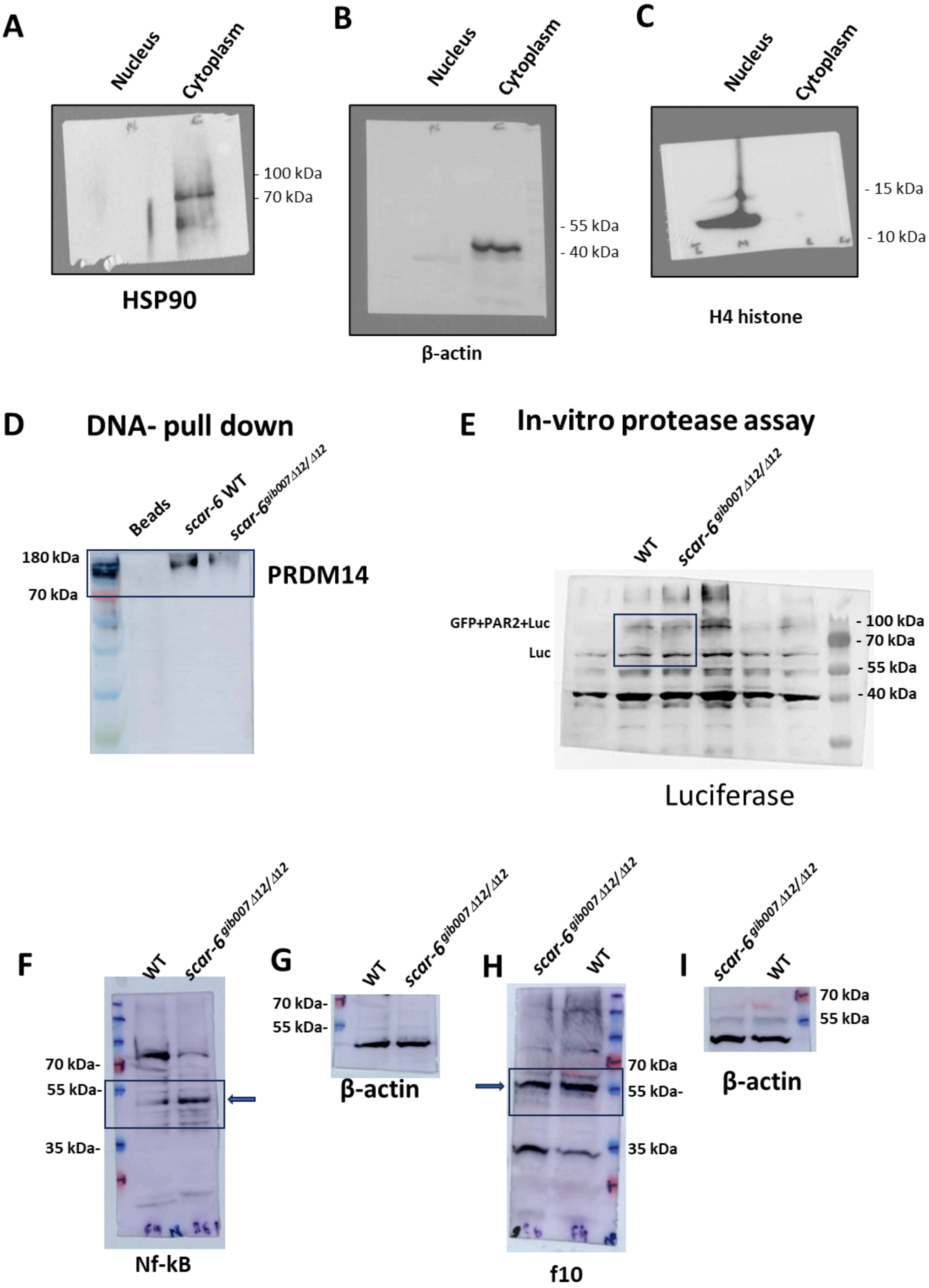
A
Nucleus
Cytoplasm
- 100 kDa
- 70 kDa
HSP90
B
Nucleus
Cytoplasm
- 55 kDa
- 40 kDa
β-actin
C
Nucleus
Cytoplasm
- 15 kDa
- 10 kDa
H4 histone
D
DNA- pull down
Beads
scar-6 WT
scar-6 gib007 Δ12/Δ12
180 kDa
70 kDa
PRDM14
E
In-vitro protease assay
WT
scar-6 gib007 Δ12/Δ12
GFP+PAR2+Luc
Luc
- 100 kDa
- 70 kDa
- 55 kDa
- 40 kDa
Luciferase
F
WT
scar-6 gib007 Δ12/Δ12
70 kDa-
55 kDa-
35 kDa-
Nf-kB
G
WT
scar-6 gib007 Δ12/Δ12
70 kDa-
55 kDa-
β-actin
H
scar-6 gib007 Δ12/Δ12
WT
70 kDa
55 kDa-
35 kDa
f10
I
scar-6 gib007 Δ12/Δ12
WT
70 kDa
55 kDa
β-actin

**Supplementary Figures 17 : -Raw images for western blot**

[A, B, C] Sub-cellular fractionation assay probing for HSP90, β-actin and H4 histone antibody respectively

[D] DNA pulldown experiment of *scar-6* locus in wild type and *scar-6*$^{gib007\Delta12/\Delta12}$ mutant DNA probing with *PRDM14* antibody.

[E] In-vitro protease assay probing with luciferase antibody.

[F-G] Western blot in wildtype and *scar-6*$^{gib007\Delta12/\Delta12}$ mutant zebrafish for Nf-κB and β-actin.

[H-I] Western blot in wildtype and *scar-6*$^{gib007\Delta12/\Delta12}$ mutant zebrafish for *f10* and β-actin.
